# Combinatorial Microgels for 3D ECM Screening and Heterogeneous Microenvironmental Culture of Primary Human Hepatic Stellate Cells

**DOI:** 10.1101/2023.05.05.539608

**Authors:** Hyeon Ryoo, Gregory H. Underhill

## Abstract

Non-alcoholic fatty liver disease affects 30% of the United States population and its progression can lead to non-alcoholic steatohepatitis (NASH), which can result in cirrhosis and hepatocellular carcinoma. NASH is characterized by a highly heterogeneous liver microenvironment created by the fibrotic activity of hepatic stellate cells (HSCs). While HSCs have been widely studied in 2D, further advancements in physiologically-relevant 3D culture platforms for the in vitro modeling of these heterogeneous environments are needed. In this study, we have demonstrated the use of stiffness-variable, ECM protein-conjugated polyethylene glycol microgels as 3D cell culture scaffolds to modulate HSC activation. We further employed these microgels as a high throughput ECM screening system to identify HSC matrix remodeling and metabolic activities in distinct heterogeneous microenvironmental conditions. In particular, 6 kPa fibronectin microgels were shown to significantly increase HSC matrix remodeling and metabolic activities in single or multiple component microenvironments. Overall, heterogeneous microenvironments consisting of multiple distinct ECM microgels promoted a decrease in HSC matrix remodeling and metabolic activities compared to homogeneous microenvironments. We envision this ECM screening platform being adapted to a broad number of cell types to aid the identification of ECM microenvironments that best recapitulate the desired phenotype, differentiation, or drug efficacy.

## INTRODUCTION

Non-alcoholic fatty liver disease (NAFLD) is the leading cause of liver-related morbidity and mortality worldwide, with an estimated 30% of people in the United States being affected by it.^[1]^ Non-alcoholic steatohepatitis (NASH) is NAFLD’s more aggressive subtype that can present itself in more than half of patients with NAFLD.^[2]^ NASH is characterized by cell injury, inflammatory cell infiltration and hepatocyte swelling that may further progress to fibrosis, cirrhosis and hepatocellular carcinoma.^[3]^ Notably, NASH has been suggested to promote a highly heterogeneous fibrotic phenotype, in comparison to other causes such as HCV.^[4]^ Hepatic stellate cells (HSCs) are now established as the main effector of liver fibrosis.^[5]^ HSCs reside in the perisinusoidal space, in close contact to hepatocytes, endothelial cells and nerve endings.^[6]^ Liver injury through causes such as viral infection, alcoholic liver disease, and NASH can lead to the activation of HSCs, making them highly proliferative and fibrogenic, leading to the accumulation of extracellular matrix and thus fibrosis.^[5,7]^ HSCs are known to interact heavily with its surrounding ECM for the regulation of its activation fate.^[8–11]^ For example, a 2D high-throughput study of substrate stiffness and ECM composition revealed that pre-activated HSCs expressed higher levels of fibrogenic proteins in softer substrates, with collagen III and IV increasing their proliferation.^[12]^ Yet, the interaction of HSCs and its ECM in the more *in vivo*-relevant 3D context is largely unexplored.

3D cell culture platforms, in most contexts, are a better representation of *in vivo* environments, due to its lack of prescribed polarity and the availability of adhesion sites in all three dimensions, leading to more relevant morphologies and behavior.^[13,14]^ Synthetic materials such as polyethylene glycol (PEG) are popular as a 3D encapsulation platform due to their highly tunable yet inherently bioinert nature. These properties allow for the independent control of the relevant cues being studied, such as adhesion ligand presence and its density, degradability, viscoelasticity and stiffness, as these can be added on to the “blank state” material as needed.^[15,16]^ The major drawback of PEG hydrogels is its nanoporous nature, as it’s highly restrictive of cell growth, it prevents free diffusion of larger molecules and makes analysis methods such as immunostaining difficult, even in the presence of proteolytically degradable crosslinkers.^[17–20]^ To circumvent this drawback, multiple studies have attempted to create a microporous PEG hydrogel, but these are often multi-step processes that disqualify their application as a high throughput system.^[21–24]^ Microgel (µG) scaffold technologies can overcome both of these drawbacks as they enable the formation of transferable, pre-formed scaffold units that are modular, heterogeneous and whose assembly can create microporous structures.^[25–28]^

Here, we introduce the use of ECM protein-tagged PEG microgels of variable stiffnesses as 3D scaffolds for the culture of *in vitro* passaged (activated) primary human HSCs. The study of HSCs in 3D has been primarily limited to spheroid^[29,30]^ or decellularized liver ECM,^[31,32]^ where it is difficult to isolate the microenvironmental factors affecting cell behavior. PEG µG scaffolds allow us to isolate these factors by the selective modification of either the stiffness or the protein composition of the µGs. After the culture of HSCs in homogeneous, single-component scaffolds, we explore the mixing of the different µGs as building blocks for the formation of a combinatorial scaffold with discrete microenvironmental cues, serving as an *in vitro* model of the highly heterogeneous microenvironment of the NASH fibrotic liver.^[4]^ While µG interparticle heterogeneity is often considered one of its main strengths,^[25]^ this property has been used only in low throughput contexts and has not been widely studied.^[33,34]^ Additionally, despite the increasing understanding of the importance of the ECM on cellular behavior, the majority of 3D high throughput studies primarily focus on the variation of the cell culture medium,^[35–38]^ with only a small number of studies modifying the encapsulating ECM.^[39,40]^ Here, we introduce the use of protein-tagged microgels of variable stiffnesses as models of 3D heterogeneous microenvironments and the exploitation of their combinatorial potential to assess HSC matrix remodeling and metabolic activity in a high throughput ECM screening system.

## RESULTS

### Microgel Composition and Characterization

The µGs employed here were fabricated by first covalently thiolating the proteins of interest using a bifunctional PEG linker, SVA-PEG-SH. The SVA reacts with the free lysines in the protein of interest to effectively convert the free amines into free thiols. The thiolated proteins were then added in a solution with multi-arm PEG norbornene, LAP (a photoinitiator), and PEG dithiol at a slight molar deficiency compared to the available norbornenes. This step served to allow the full integration of the thiolated proteins and to reduce the competition of the crosslinking PEG dithiol. 24% dextran and 16% MgSO4 solution was added to this PEG solution, creating an aqueous two-phase system that can be vortexed to form a microdroplet suspension of PEG within the dextran bath.^[41,42]^ The suspension was irradiated with UV light for 90 seconds to allow for the microdroplets to polymerize and form a hydrogel **(Figure 1a)**. Interestingly, when the protein was covalently crosslinked to the PEG backbone at the same time as the polymerization, the protein was distributed in the core or on the outer edge of the µG. Meanwhile if the protein was pre-crosslinked to the PEG backbone before the polymerization, the protein was evenly distributed within the µG, possibly due to an increase in hydrophilicity of the proteins **(Supplementary Figure 1)**. Based on the goal of promoting surface presentation of ECM proteins and subsequent interactions with cells, subsequent studies incorporated the simultaneous protein crosslinking and gel polymerization.

**Figure 1.**
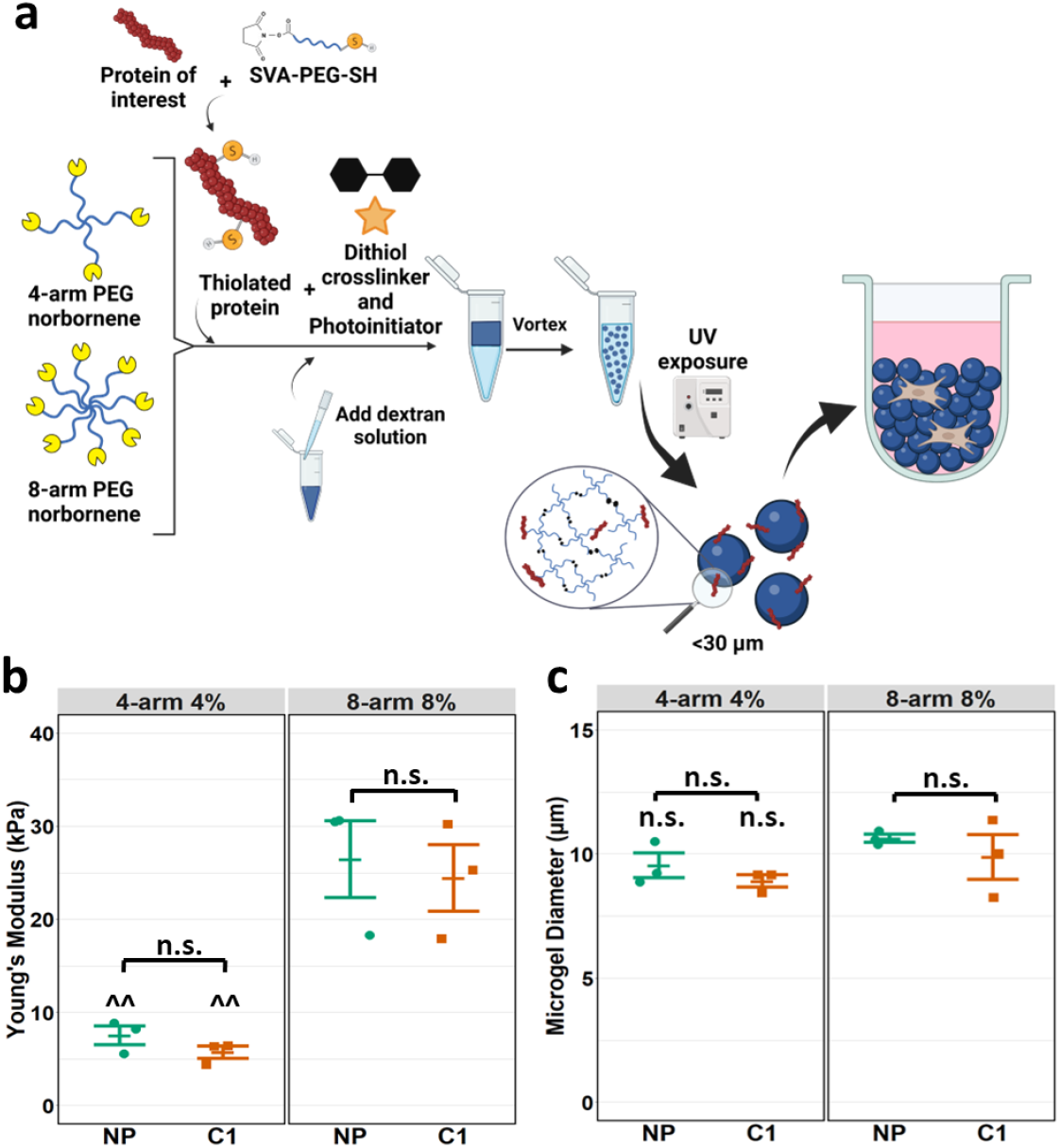
**a)** Diagram of 4-arm 4% or 8-arm 8% microgel production. The protein of interest is functionalized with thiols using SVA-PEG-SH. The thiolated protein is added to the multi-arm PEG along with PEG-dithiol and LAP photoinitiator. Dextran/MgSO4 solution is added to the PEG, making an aqueous two-phase system. This system when vortexed, forms PEG microdroplets that can be polymerized with UV exposure. The microgels are then filtered through a 30 µm membrane. These can then be added with cells in a confined system as a 3D scaffold. **b)** 4-arm 4% (4) and 8-arm 8% (8), no protein (NP) and collagen I-conjugated (C1) µGs’ Young’s modulus. n=3 experimental replicates of 10 µGs each. **c)** 4NP, 4C1, 8NP and 8C1 µGs’ diameter. n=3 experimental replicates of 300 µGs each. Average ± standard error of means. Two-way interaction ANOVA analysis. n.s. p > 0.05 and ^^ p < 0.01 vs 8-arm 8% counterpart.

We utilized atomic force microscopy (AFM) to determine µG stiffness. 4-arm PEG-norbornene at 4% w/v µG composition corresponded to a Young’s modulus of 7.55 ± 1.01 kPa. For the stiffer µGs, 8-arm PEG-norbornene was composed to 8% w/v (8NP) resulting in a Young’s modulus of 26.45 ± 4.10 kPa (Figure 1b). To confirm that the µGs retain their stiffness when proteins are incorporated into the µG, collagen I tagged µGs were measured. 4-arm 4% µGs with collagen I (4C1) had an elastic modulus of 5.72 ± 0.68 kPa and the 8-arm 8% µGs with collagen I (8C1) had an elastic modulus of 24.44 ± 3.58 kPa **(Figure 1b)**. While the stiffness of the µGs appeared to modestly decrease with the addition of collagen I, no significant differences were observed **(Figure 1b)**. This range of stiffnesses is physiologically relevant to the progression of liver fibrosis. Specifically, the 4-arm 4% µG stiffness (6-7 kPa) is typically associated with the early/intermediate phase of liver fibrosis, while 8-arm 8% gels (24-26 kPa) exhibit the stiffness associated with advanced fibrosis.^[12,43]^ 4NP and 4C1 had µG sizes of 9.53 ± 0.50 µm and 8.91 ± 0.24 µm respectively. 8NP and 8C1 had µG sizes of 10.63 ± 0.16 µm and 9.88 ± 0.90 µm respectively **(Figure 1c)**. No significant differences in µG diameter between the different PEG composition or collagen I presence conditions were observed **(Figure 1c)**. This small size of the µGs was desired, as it would enable cultured cells to interact with more than just a few µGs at a time, emulating the heterogeneous environment of fibrotic livers while allowing for the study of the combinatorial effect of these on the cells.

### Microgel Protein Content Modulates HSC 3D Morphology

Having observed the integration of proteins into the µG as well as established variable stiffness µG compositions, µGs were tested as 3D culture scaffolds for activated primary human HSCs. µGs of both 4-arm PEG 4% (4) and 8-arm PEG 8% (8) compositions were conjugated with the ECM proteins fibronectin (FN), collagen I (C1), collagen III (C3), collagen IV (C4) or with no protein (NP). As expected, HSCs cultured with both 4NP and 8NP conditions preferred to aggregate with each other over interacting with the non-fouling PEG µGs **(Figure 2a)**. In contrast, all ECM protein-conjugated µGs enabled HSC attachment at different extents **(Figure 2a)**. For the quantification of morphological factors such as cell spreading and cell aggregation, the minimum distance between nuclei was measured for each nucleus and averaged per 3D image. 4NP and 8NP µGs had significantly lower distance between nuclei, indicating high cell aggregation **(Figure 2b)**. For the µGs with ECM protein attachment, we observed no significant differences in the distance between the nuclei for 4-arm 4% µGs but observed that 8FN and 8C4 µGs had significantly higher distance between nuclei with 39.88 and 37.26 µm respectively, compared to 8C1 or 8C3 µGs with 27.57 and 24.07 µm respectively **(Figure 2c)**. Also, HSCs cultured in 4FN (31.01 µm) had significantly lower and 4C3 (31.37 µm) had significantly higher distance between nuclei when compared to their 8-arm 8% counterparts **(Figure 2c)**. This result implied that while ECM protein effects on cell aggregation or migration are minor when the stiffness of the scaffold is in the lower range, when exposed to a higher stiffness, HSCs demonstrate a more pronounced spreading behavior that is regulated by the type of ECM protein present. Comparably, a seminal study on the effect of stiffness and ECM protein on HSC activation had previously shown that the differences in cell spreading caused by ECM proteins were more pronounced on HSCs cultured in stiff substrates over softer ones.^[8]^

**Figure 2.**
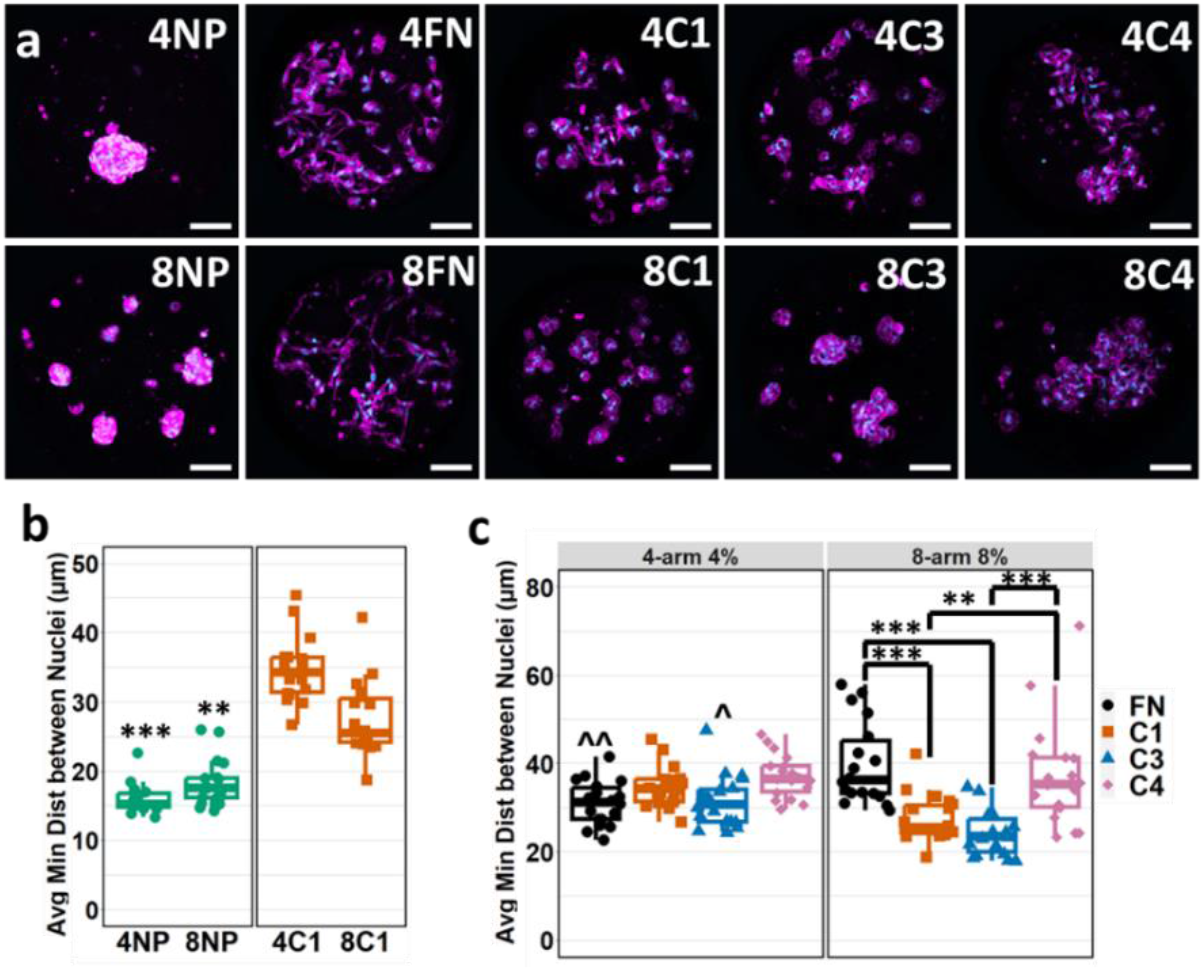
**a)** Maximum intensity projection of confocal images of HSCs cultured in PEG microwells along with their respective µGs of 4-arm 4% (4) and 8-arm 8% (8) compositions and no protein (NP), fibronectin (FN), collagen I (C1), collagen III (C3) and collagen IV (C4) conjugation. Purple: DiD membrane stain. Light blue: DAPI. Scale bar: 100 µm. **b)** Box and whisker plot of the average minimum distance between nuclei of all HSC nuclei within each microwell of 4NP, 8NP, 4C1 and 8C1 scaffolds. n = 15-18 from 5-6 experimental replicates. Two-way interaction ANOVA. *** p < 0.001 vs C1 counterpart. **c)** Box and whisker plot of the average minimum distance between nuclei of all HSC nuclei within each microwell of ECM protein-conjugated µG scaffolds. n = 18 from 6 experimental replicates. Two-way interaction ANOVA. * p < 0.05 ** p < 0.01 *** p < 0.001 where * means between conditions and ^ means vs 8-arm 8% counterpart.

### Microgels Modulate HSC Protein and Gene Expression

To assess the effect of µG composition on HSC activation, we performed immunocytochemistry and RT-qPCR on known HSC activation markers. Immunostaining experiments of HSCs revealed significant differences in the expression of platelet derived growth factor receptor beta (PDGFRβ) and collagen I, which was dependent on the ECM and stiffness of the µGs present in the culture **(Figure 3)**. HSCs cultured with 8FN had significantly higher PDGFRβ expression (1.32 AU) compared to HSCs cultured in other 8-arm 8% µGs while 8C4 showed the lowest expression of PDGFRβ (0.60 AU) **(Figure 3b)**. In contrast, 8C4 showed the highest expression of collagen I, with a fluorescence 1.57 times the average of all conditions tested **(Figure 3d)**. This contrasting expression of both activation markers on HSCs cultured in 8C4 scaffolds indicated that the phenotypic states of HSCs could be more nuanced than mere high/low activation. HSCs also showed differential expression dependent on the µGs stiffness. For example, HSCs on 8C4 µG scaffold showed significantly lower PDGFRβ and significantly higher collagen I expression compared to HSCs on 4C4 µG scaffold **(Figure 3b,d)**. Two-way interaction ANOVA also revealed higher stiffness µGs tended to decrease PDGFRβ expression while increasing collagen I expression **(Figure 3b,d)**. Collagen I-tagged µGs decreased lysyl oxidase expression in comparison to other protein-tagged µGs **(Supplementary 2a)**, while no significant differences were observed between the HSCs cultured in µG scaffolds for α smooth muscle actin (αSMA) expression **(Supplementary 2b)**. Once again, the exposure of HSCs to a greater stiffness scaffold led to an increasing distinction on how the cells behaved within the different ECM protein scaffolds, implying a combinatorial effect between ECM protein and stiffness that are not merely additive.

**Figure 3.**
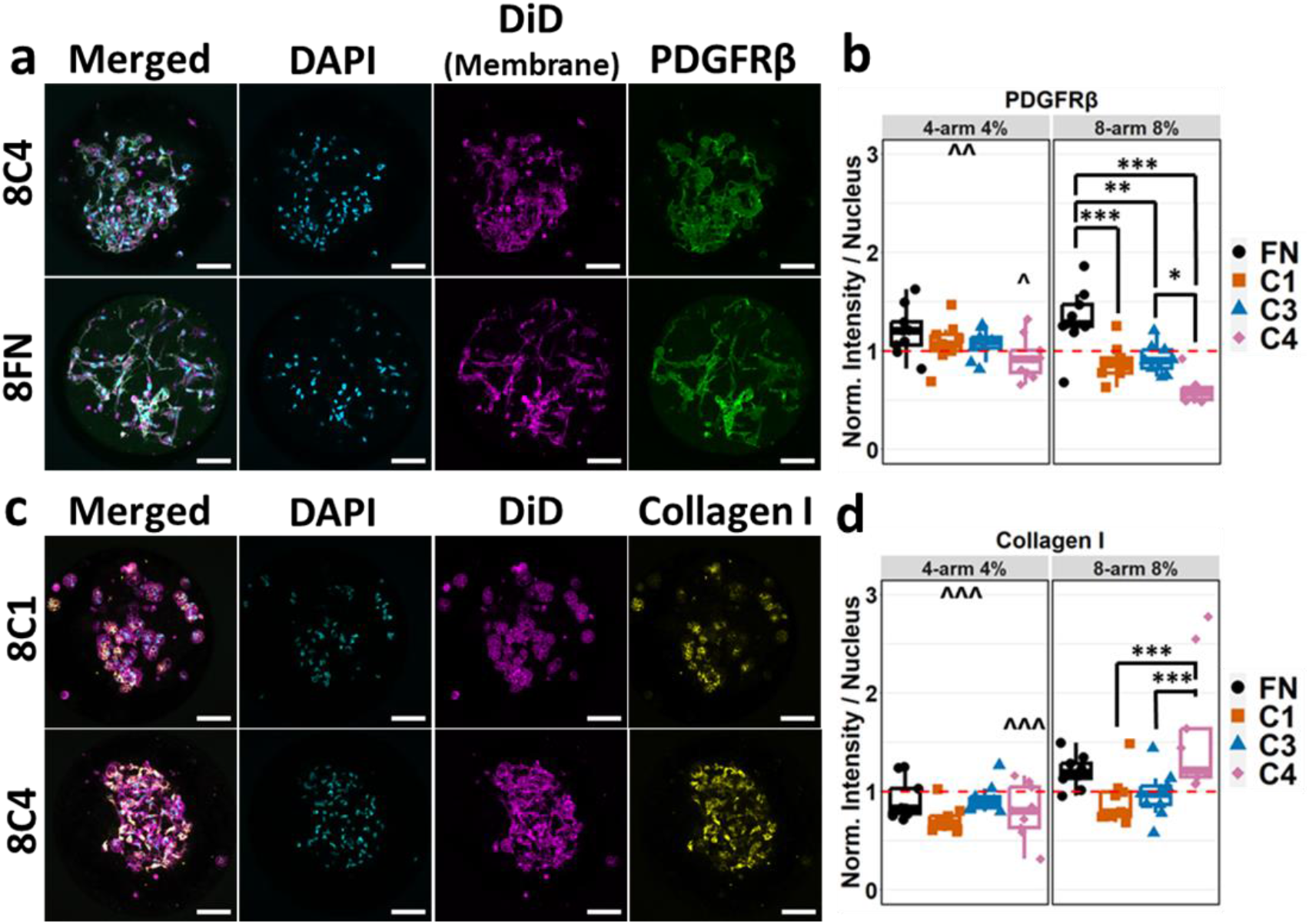
**a)** Representative maximum intensity projection of confocal images of HSCs cultured with 8C4 and 8FN scaffolds, the lowest and highest conditions for PDGFRβ expression. Light blue: DAPI. Purple: DiD membrane stain. Green: anti-PDGFRβ. Scale bar: 100 µm. **b)** Box and whisker plots of the anti-PDGFRβ fluorescence intensity per nucleus normalized to the average of all conditions per experimental replicate. n = 9 from 3 experimental replicates. **c)** Representative maximum intensity projection of confocal images of HSCs cultured with 8C1 and 8C4 scaffolds, the lowest and highest conditions for collagen I expression. Light blue: DAPI. Purple: DiD membrane stain. Yellow: anti-collagen I. Scale bar: 100 µm. **d)** Box and whisker plots of the anti-collagen I fluorescence intensity per nucleus normalized to the average of all conditions per experimental replicate. n = 9 from 3 experimental replicates. Two-way interaction ANOVA. * p < 0.05 ** p < 0.01 *** p < 0.001 where * means between conditions and ^ means vs 8-arm 8% counterpart.

RT-qPCR was performed on 9 different activation marker genes of HSCs. In average, all 9 genes showed elevated expression when cultured in 8-arm 8% scaffolds vs 4-arm 4% scaffolds. When testing for significance, two-way interaction ANOVA analysis revealed that 6 of the 9 tested genes had significantly higher expression in the stiffer scaffolds **(Figure 4a)**. Performing a principal component analysis (PCA) on the qPCR dataset separated the 8 conditions into three different phenotypic clusters. The top two principal components explained 85.84% of the variance in the data. Cluster 1, composed of 4C1, 8FN, 4FN and 4C3, was characterized by low MMP2, IL6 and ACTA2. Cluster 2, composed of 8C4, 8C1 and 4C4, was characterized by high CDH2 and COL1A1. Cluster 3 was composed solely of 8C3 and was characterized by high TIMP1, TGFB1 and ACTA2 **(Figure 4b)**. This result was indicative of the different phenotypes that could arise from the activated HSCs being cultured in different microenvironments. A previous 2D study on activated HSCs had also identified HSCs with different HSC activation phenotypes.^[12]^ Similarly, *in vivo* single cell RNAseq studies have shown activated HSC phenotypic variability within fibrotic mouse livers.^[44,45]^

**Figure 4.**
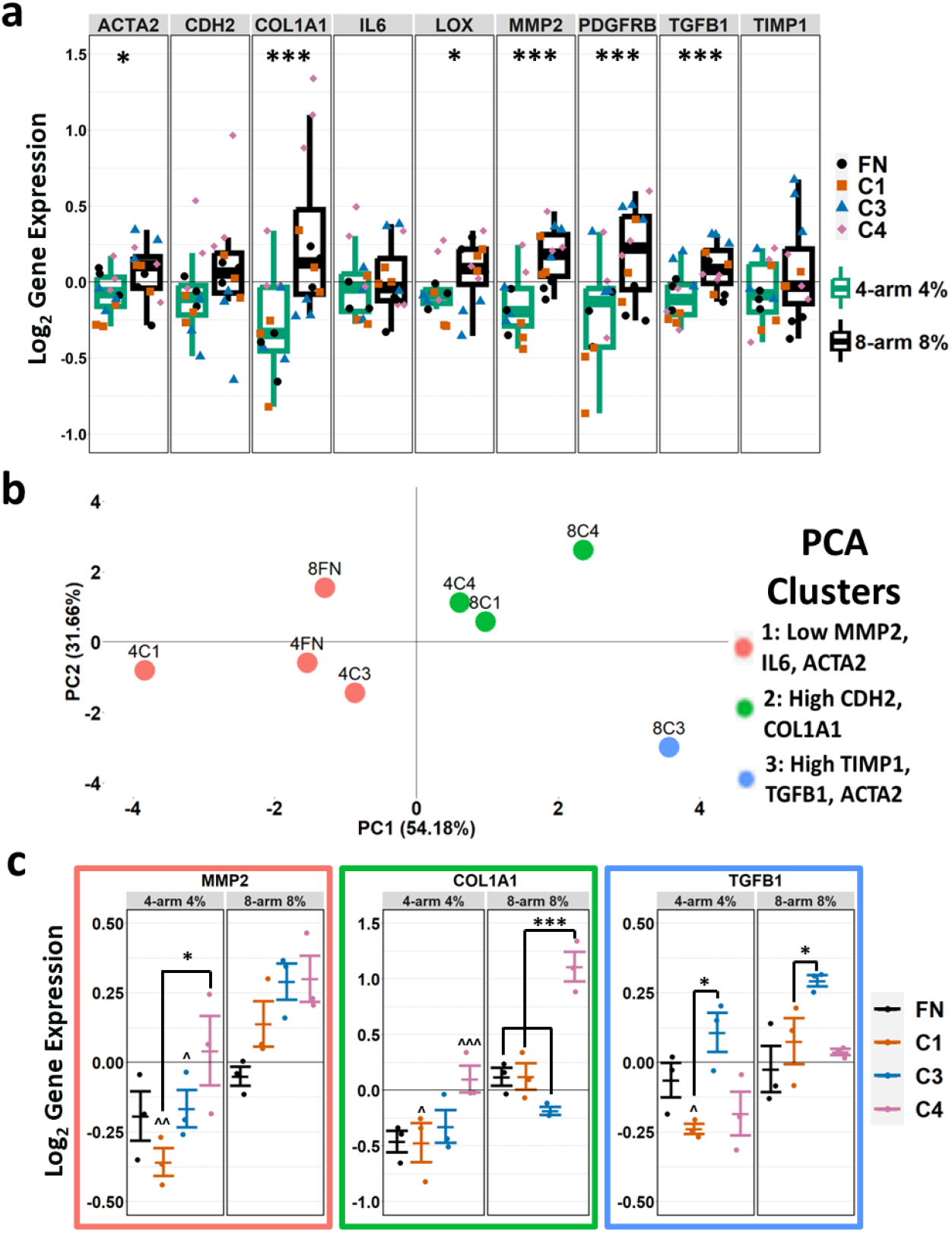
**a)** Box and whisker plot of the 9-gene RT-qPCR data on HSCs cultured with 8 conditions of µGs, separated primarily by the PEG backbone composition. Each dot represents an experimental replicate of HSCs cultured in the corresponding ECM protein conjugation condition within the PEG backbone composition. n = 3 experimental replicates for each multi-arm PEG composition and ECM protein combination. Two-way interaction ANOVA analysis.* p < 0.05 *** p < 0.001 between 4-arm 4% and 8-arm 8% conditions. **b)** Plot of the top two principal components of the PCA performed on the RT-qPCR data. **c)** Average ± standard error of means for three of the genes highlighted in the PCA analysis. n = 3 experimental replicates. Two-way interaction ANOVA analysis. * p < 0.05 ** p < 0.01 *** p < 0.001 where * means between conditions and ^ against its 8-arm 8% counterpart.

Focusing in on the representative genes of each cluster revealed the clear distinctions between each cluster’s leading condition (4C1 for cluster 1, 8C4 for cluster 2 and 8C3 for cluster 3) and the rest of the conditions tested **(Figure 4c)**. It is interesting to note the phenotypic distinction of HSCs in these microenvironments in which ECM and stiffness can be independently controlled. In previous studies, collagen I has been demonstrated to be highly abundant in fibrotic livers, thus stiffer livers, while collagen IV levels are downregulated or remain similar in fibrosis, and thus are proportionately higher in soft, healthy livers.^[5,46,47]^ Our results indicate that ECM and stiffness can influence HSC phenotype in a combinatorial manner, suggesting that potential spatial or temporal variations in the evolution of ECM and stiffness changes in the liver could likely promote a complex array of phenotypic responses. Further, the gene expression dataset also revealed additive and non-additive behavior depending on the stiffness of the scaffold or the ECM protein content. As one example, COL1A1 expression independently increased with the conjugation of collagen IV and the use of 8-arm 8%, with the combination of these two factors synergistically elevating COL1A1 expression **(Figure 4c)**. Conversely, while LOX expression tended to increase with the use of the stiffer µGs, HSCs in C3 µGs showed lower LOX expression in 8C3 µGs compared to 4C3 µGs **(Supplementary Figure 3)**. While the identification of general trends of HSC behavior as an effect of stiffness or ECM protein can be valuable, this type of non-additive or synergistic behavior justifies the need for empirical tests of each ECM condition in studying HSC activation phenotype.

### High Throughput Combinatorial 3D ECM Screening

For the high throughput combinatorial platform, the same 8 types of µGs (4FN, 4C1, 4C3, 4C4, 8FN, 8C1, 8C3, 8C4) were used. By employing a liquid handler, these µGs were distributed at varying ratios across different wells of a 384-well plate. Duplicate wells of single component, double component (half-half) and triple component (third-third-third) µGs were produced. After two days of culture, the plates were added with either MMP2 substrate or resazurin to measure MMP2 activity or metabolic activity using a plate reader **(Figure 5a,b)**. The average of 3 experiments of MMP2 readout and 3 experiments of resazurin readout for each µG composition condition were analyzed and plotted in Figure 5c. MMP2 (matrix metalloproteinase-2) is a matrix remodeling protein that becomes overexpressed in activated HSCs and serum MMP2 levels could even serve as good indicators of liver cirrhosis.^[48,49]^ MMP2 can exist in a pro- and active form and this conversion is tightly regulated by TIMP (tissue inhibitor of metalloproteinase) and MT1-MMP (membrane-type 1 metalloproteinase) proteins, all highly regulated by HSCs and thus MMP2 activity is an indication of the HSCs’ proneness to matrix remodeling.^[48]^ Resazurin is a molecule that becomes reduced into resorufin under active metabolism of viable cells.^[50]^ Resorufin is highly fluorescent and can serve as a sensitive readout of cells’ metabolic activity, which can also correlate to cell viability, proliferation and survival. We employed these two readouts for their clear relevance in HSC function and activation state, as well as their compatibility with plate reader platforms and thus suitability for use in high throughput experiments.

**Figure 5.**
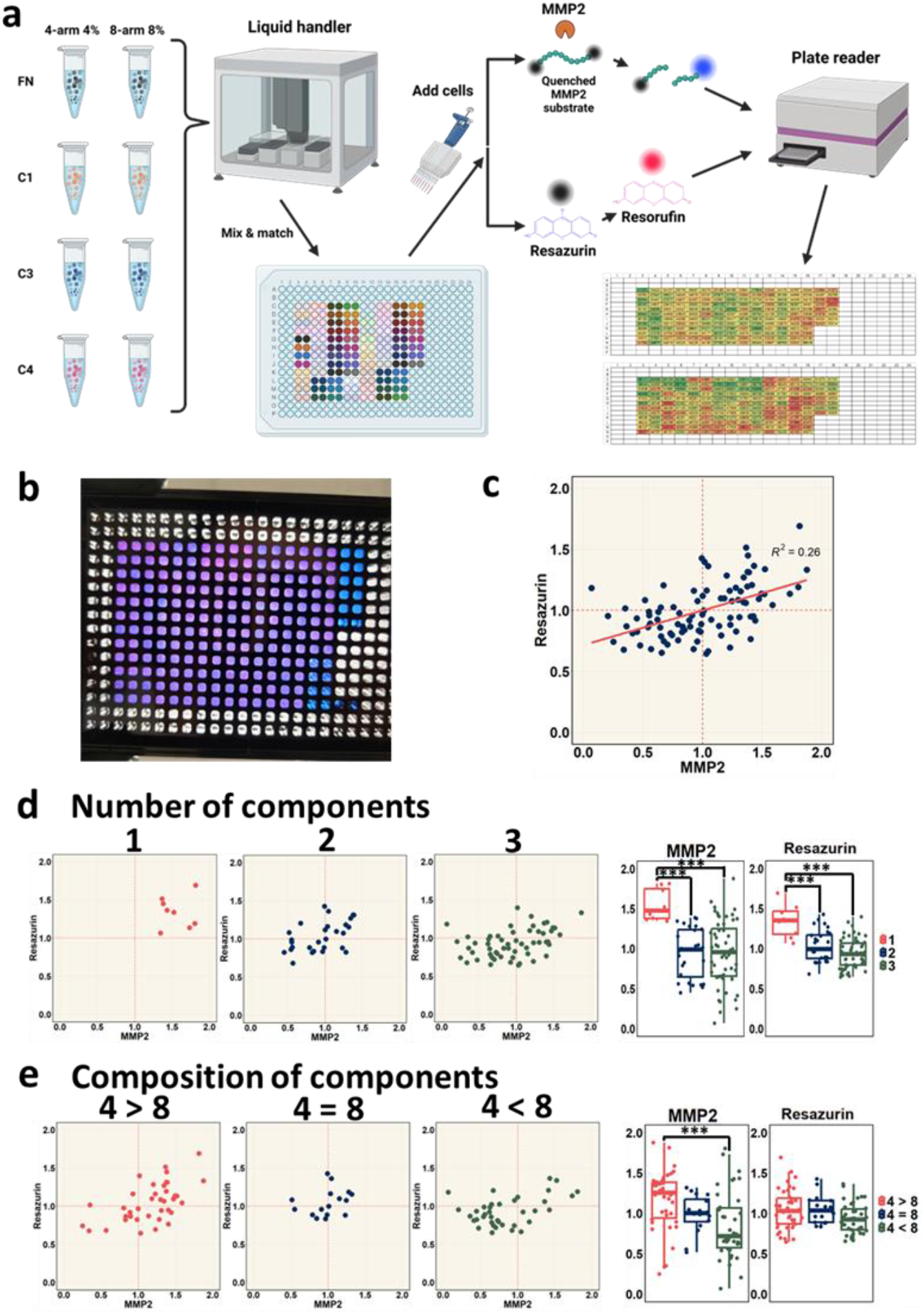
**a)** Diagram of the creation and analysis of the high throughput ECM screening platforms. Pre-formed µGs of different stiffness or ECM protein are distributed in predetermined ratios using a liquid handler. Cells are then added onto the plate by using a multichannel pipettor or a liquid handler and cultured for 2 days. For analysis, fluorescent or chromogenic substrates such as MMP2 degradable fluorescence-quenched substrates or resazurin are added into each well and measured in a plate reader. **b)** Photograph of a 384-well plate prepared for high throughput ECM screening, after 2 hr incubation in resazurin. **c)** Scatter plot of the n=3 experimental replicate average of normalized MMP2 and resazurin values, with an R squared correlation value of 0.26. Values are normalized to the average of all conditions per experimental replicate and then averaged across experimental replicates to produce one data point per condition. **d)** Subset scatter plots of Figure 5c, separated by the number of components in each condition, with quantification of the distribution using box and whisker plots. **e)** Subset scatter plots of Figure 5c, separated by the composition of each condition, with quantification of the distribution using box and whisker plots.. 4 > 8 means there’s more 4-arm 4% μGs than 8-arm 8% μGs within the condition, disregarding the ECM protein composition. 4 = 8 means there’s equal ratios of the two types of gels. 4 < 8 means there’s more 8-arm 8% μGs than 4-arm 4% μGs within the condition. One-way ANOVA analysis. *** p < 0.001.

The HSCs’ MMP2 and metabolic activity had a weak correlation with an R squared value of 0.26 **(Figure 5c)**. Interestingly, the number of components in the scaffold significantly affected both MMP2 and resazurin readouts. Single component conditions had an average normalized MMP2 activity of 1.55 AU compared to double component of triple component conditions with 0.94 and 0.95 AU respectively. For metabolic activity, single component conditions had an average of 1.34 AU while double component conditions had an average of 1.03 AU and triple component conditions had an average of 0.93 AU **(Figure 5d)**. This implied that the heterogeneity of the scaffold could negatively regulate the HSCs’ MMP2 and metabolic activity. For example, 4FN and 8C3 ranked as 2^nd^ and 4^th^ highest MMP2 readouts with 1.81 and 1.73 AU, but this decreased to 1.05 AU in 4FN+8C3 scaffolds. Similarly for resazurin, 4C4 and 4C1 had readouts of 1.51 and 1.45 AU but it had a reduced value of 0.82 AU when mixed into one scaffold. Another broad trend we could observe was the effect that the composition of the component µGs had on the two readouts. Meaning, when the scaffold had mostly softer gels compared to stiffer gels (4 > 8), the HSCs behaved differently to when the scaffold had mostly stiffer gels (4 < 8). HSCs cultured on scaffolds with primarily 4-arm 4% µGs had an MMP2 activity of 1.16 AU, while HSCs cultured on scaffold with primarily 8-arm 8% µGs or the same ratio (4 = 8) between the two had an MMP2 activity of 0.84 and 0.99 AU respectively **(Figure 5e)**. This effect was most clear between the 4FN+4C1+4C4 and 8FN+8C1+8C4 conditions, as 4FN+4C1+4C4 scaffolds had the highest MMP2 activity with 1.88 AU but the higher stiffness 8FN+8C1+8C4 scaffold had an MMP2 readout of 0.62 AU.

To delve deeper into the effect each individual type of µG had on the scaffold, heatmaps of the different conditions were generated along with linear regression analysis and ranked 95% confidence interval plots **(Supplementary Figure 4)**. All linear regression coefficients are given as a comparison against the 8FN component, or 8FN as the intercept, which was typically the component with the median coefficient value. For single component conditions, both MMP2 and metabolic activity readouts were highest on 4FN scaffolds with 1.81 and 1.69 AU respectively. 4C3 had the lowest MMP2 and metabolic activity readouts with 1.35 and 1.07 AU respectively **(Figure 6a)**. Double component conditions showed similar trends as the single component conditions, with conditions that included 4FN showing higher MMP2 and metabolic activity readouts compared to conditions without **(Figure 6b)**. As expected, the linear regression analysis of only the conditions with double components revealed that 4FN had the greatest increasing effect on both MMP2 and resazurin readouts, with a linear coefficient of +0.72 and +0.60 respectively when compared against the 8FN component. In contrast, 8C1 had the lowest MMP2 coefficient with -0.41 and 4C1 had the lowest resazurin coefficient with -0.21 **(Supplementary Figure 5a**,**b)**. For triple component conditions, the linear regression revealed the components that lowered the readouts the most were 8C3 for MMP2 and 8C1 for resazurin readouts, **(Supplementary Figure 5c**,**d)** thus these subset data were selected for demonstration in **Figure 6c**. While throughout all conditions 4FN was an increasing factor for both readouts, its effect was greatest in the triple component conditions, as its regression coefficient reached 1.81 for MMP2 readouts and 0.83 for resazurin readouts vs 8FN baseline **(Supplementary Figure 5c**,**d)**. This trend was clear to see in the 3-component raw value heatmaps too, as conditions with 4FN as a component showed consistently high values (average of 1.28 AU for MMP2 and 1.10 AU for resazurin) while the conditions with 8C3 and 8C1 as components had low MMP2 or resazurin readouts with an average of 0.83 AU and 0.86 AU respectively **(Figure 6c)**.

**Figure 6.**
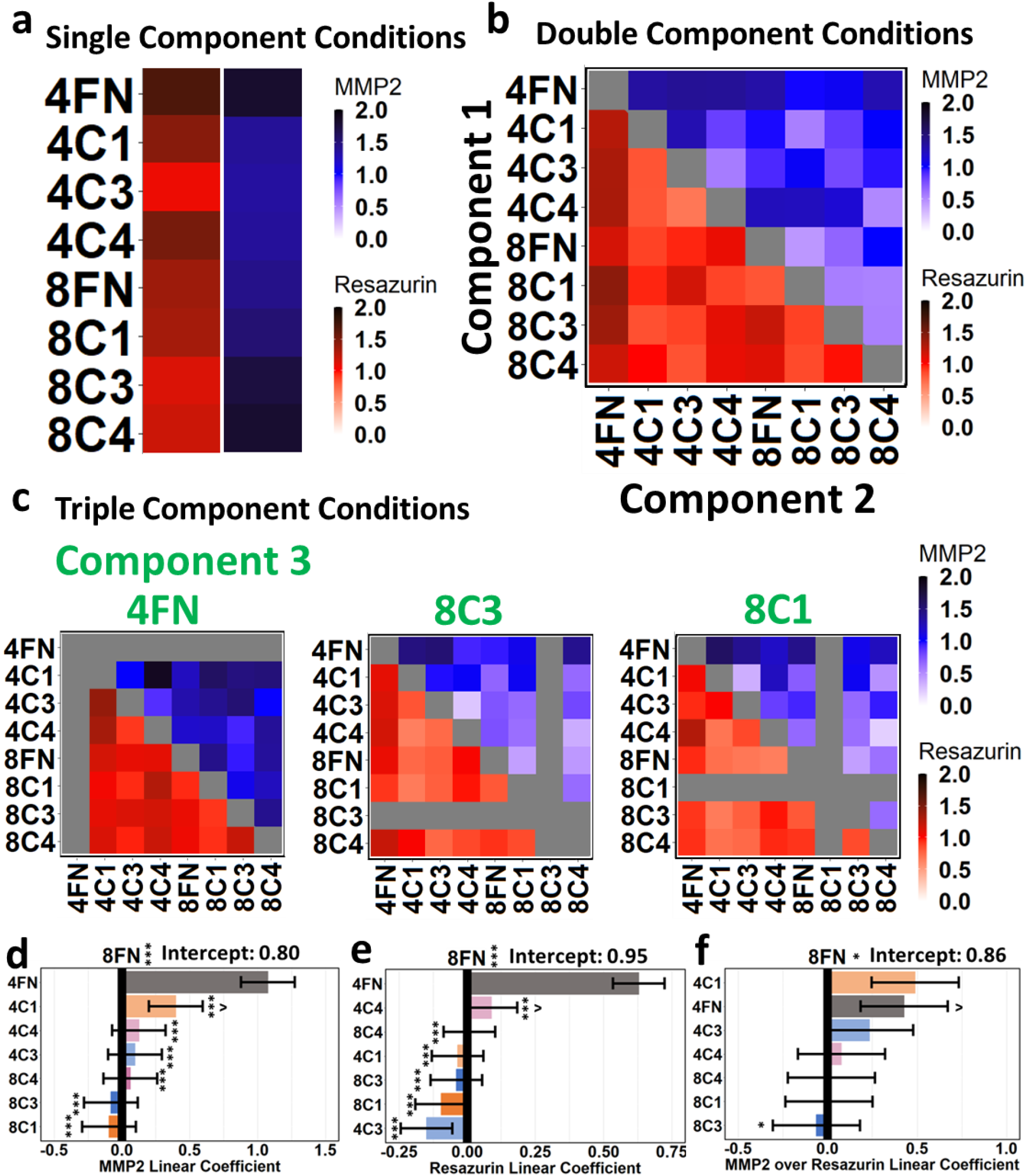
**a)** MMP2 and resazurin readout heatmaps for conditions in which only one component was present. **b)** MMP2 and resazurin readout heatmaps for conditions in which two components were present. **c)** MMP2 and resazurin readout heatmaps for conditions in which three components were present. The components written in green are the common component between the conditions shown in each heatmap below. Subset of data was selected based on the highest (4FN) and lowest (8C3, 8C1) linear regression coefficients for MMP2 and resazurin readouts. **d**,**e**,**f)** Bar graph of linear regression coefficients for MMP2, resazurin and MMP2/resazurin readouts with 8FN as the intercept. F-statistic on linear regression model. Error bars indicate standard error. * p < 0.05, *** p < 0.001 with * meaning vs highest condition (4FN, 4FN, 4C1) and ^ meaning vs lowest condition (8C1, 4C3, 8C3).

When all 1, 2 and 3 component conditions are accounted for, 4FN remained the component that positively influenced MMP2 and metabolic activity most profoundly, while 8C1 was the most negative influencer of MMP2 activity and 4C3 was the most negative influencer of metabolic activity **(Figure 6d,e)**. Interestingly, when the average of the MMP2 measurements for each condition were normalized by the average of the resazurin measurements for each condition and subjected to linear regression analysis, 4FN did not have the most positive linear regression coefficient and was surpassed by the 4C1 component **(Figure 6f)**. This indicated that 4FN components led to a relatively proportional increase of MMP2 and metabolic activity, while 4C1 components might be more impactful specifically towards MMP2 activity. Also of interest were conditions with triple components that greatly varied its MMP2 activity while retaining a similar metabolic activity. For example, condition 4C4+8C1+8C4 had one of the lowest MMP2 activities with 0.20 AU, while a very similar condition 4C1+4C4+8C4 demonstrated high MMP2 activity with 1.44 AU, with both resazurin readouts being similar at 0.95 and 0.92 AU respectively **(Supplementary Figure 4)**. Meaning, a small modification of only one third of its total components from a softer 4C1 to a stiffer 8C1 had led to a 7.08 fold increase in MMP2 activity. This result greatly emphasizes the importance of individual ECM components on HSC behavior and the need for ECM screening procedures in identifying the environments that modulate cell behavior in a desired way.

In this study we have explored the use of µGs as building blocks for a 3D cell culture scaffold and the exploitation of their heterogeneity potential to create a high throughput ECM screening platform in 3D. The high throughput study of cell behavior depending on ECM conditions in 3D cell culture is a largely unexplored field.^[39,40]^ Ranga et al. encapsulated cells in bulk PEG hydrogels tethered with various ECM-mimetic peptides using a liquid handler to create a 3D high throughput screening system.^[39]^ The system was the first of its category, though it relied on the use of nanoporous gels as well as relying on time-sensitive chemistries that could make liquid handling difficult. We devised this system reported here to allow for the relatively free spreading and migration of our cultured cells through the microporous structure, which also allows for the free diffusion of metabolites and analytes. It is worth recognizing that this same feature prevents this platform from tackling one of the factors studied in the fore-mentioned high throughput system, degradability-dependent cell behavior, unless interstitial matrix is also added along with the cells. Our reported system consisted of the pre-addition of modular building blocks to the plate with the use of a liquid handler, reducing the risk of affecting the cells during what could be the most stressful and time-critical moment of the cell culture, the cell seeding procedure.

Overall, we demonstrated the possibility of creating differential protein and gene expression on pre-activated primary HSCs by culturing them in scaffolds composed of 6 or 25 kPa tagged with fibronectin, collagen I, III or IV. Using this approach, it was possible to observe a trend of increasing activation marker gene expression with the use of stiffer µGs. This correlates well with the literature, as it has been shown in both 2D and 3D experiments that increasing the stiffness of the HSC substrate leads them to a more activated state.^[8,32,51]^ In future studies, it would be of interest to identify if 8-arm 8% µG bulk scaffolds have a distinct shear modulus compared to 4-arm 4% µG scaffolds and try to decouple the individual µG Young’s modulus and the bulk scaffold’s shear modulus effect on cell behavior. A related limitation of our work was that aqueous two-phase systems are less stable than oil-water emulsions and thus 4-arm PEG norbornene at 4% was the lowest concentration of PEG we could reliably produce µGs. Efforts to lower the PEG concentration and thus create healthy liver stiffness (∼1 kPa) µGs could expand upon the potential of the combinatorial µG scaffold system.

The gene expression data was also able to define three different phenotypic clusters, each with its own enhanced and diminished activation markers, further indicating the complexity of the HSC phenotype.^[44,45,52]^ High throughput studies helped identify 4FN as the most positively influential microgel in terms of MMP2 and metabolic activity across 92 different single, double and triple component microenvironments. This was especially interesting as, in a healthy liver’s perisinusoidal space, the region that the HSCs reside in, is known to be most abundant in fibronectin.^[53]^ Also, during liver injury it is one of the most upregulated ECM proteins and has been shown to be critical regulating HSC activation and survival.^[46,54,55]^ While the visualization of strong general trends was enabled through the screening of multiple different conditions, it also enabled the observation of outlier conditions such as 4C4+8C1+8C4 and 4C1+4C4+8C4, where the change of stiffness of one-third of its components led to a 7.08 fold increase in MMP2 activity while retaining a slightly lower metabolic activity. Such combinatorial regulation, including synergistic or antagonistic effects of distinct ECM proteins and stiffness have been demonstrated in previous liver-context 2D high throughput studies,^[12,56–59]^ confirming the complexity of microenvironmental interactions and making further high throughput ECM studies enticing in other organ-related contexts.

Recent studies have found that the components of the heterogeneous microenvironments of tumors could have a large effect on how the cancer cells migrate or respond to drugs.^[60,61]^ Using a 3D ECM screening system, such as the platform we report here, could be employed towards identification of ECM composition/stiffness conditions that lead to the best drug response for such cells, with the future possibility of incorporating patient-specific samples. The injectability of the µGs also means that the µGs could then be co-delivered with therapeutic agents to enhance their effect. For example, upon in vitro identification of an ECM combination that enhances the effectiveness of a drug in patient-specific tumor cells using the ECM screening system, one could deliver degradable, drug-encapsulated µGs to the site of interest. As the cells interact with the ECM-tagged µGs, they would become more receptive to the drugs being eluted from them. This could be further expanded to finding therapeutic agents for pathogenic cells that reside in other highly heterogeneous environments such as the fibrotic liver as seen here or even in homogeneous environments but have enhanced reactions to drugs in non-native ECM conditions.

## CONCLUSION

In this study, we present the creation of stiffness and protein-tunable PEG microgels and their use as combinatorial 3D cell culture scaffolds with high throughput application potential. We demonstrated that we could modulate HSC morphology, protein expression, and gene expression by providing the cells with a microporous scaffold composed of ∼6 or ∼25 kPa microgels of ∼10 μm that contained different ECM proteins such as fibronectin, collagen I, III and IV. High throughput studies revealed a trend of 6 kPa fibronectin microgels exhibiting the most profound effect on the matrix remodeling and metabolic activity of HSCs out of the 8 components tested. The high throughput system also allowed for the identification of specific ECM combinations in which small changes in the composition of the scaffold could greatly modulate the matrix remodeling and metabolic activities in the HSCs. We envision the application of this ECM screening system for the identification of ECM combinations that lead to the most desired phenotype, differentiation or drug efficiency.

## EXPERIMENTAL SECTION

### Functionalization of Proteins with Thiols

Fibronectin (FC010, MilliporeSigma), collagen I (08-115, MilliporeSigma), collagen III (M20S, Cell Guidance Systems) and collagen IV (C5533, MilliporeSigma) were prepared to 1 mg/mL by addition of deionized water if needed. 3 parts volume of 1 mg/mL protein was supplemented with 1 part volume of 1 mM SVA-PEG-SH (R-SH-0011, Ruixibio) in 1 M HEPES buffer (25060CI, Corning), mixed well and allowed to react for 4 hrs in room temperature.

### PEG Microgel Formation

All procedures were done while observing sterile practices. A 24% w/v dextran (100 kDa, 09184, MilliporeSigma) and 16% w/v MgSO_4_ (M65, Fisher Scientific) solution in PBS with 1% PenStrep (SV30010, Cytiva) (PS-PBS) was prepared by dissolving dextran and MgSO_4_ to 40% first and mixing the two at a 3:2 ratio. 4-arm PEG norbornene (20 kDa, PSB-4112, Creative PEGWorks) is dissolved to 10% w/v using 0.25% w/v LAP and 8-arm PEG norbornene (40 kDa, PSB8310, Creative PEGWorks) is dissolved to 20% w/v using 0.25% w/v lithium phenyl-2,4,6-trimethylbenzoylphosphinate (LAP, 900889, MilliporeSigma). Thiolated proteins (now 4-part volume) were mixed with 4 parts volume of the 4-arm or 8-arm PEG norbornene and 2 parts volume of 19.5 or 39 mg/mL PEG-dithiol (1 kDa, 717142, MilliporeSigma). The concentration of PEG-dithiol was chosen to allow for slightly below full occupation of the norbornenes (97.5%) as to allow for the proteins to bind to the norbornenes without competition. Dextran solution was slowly added to the PEG solution at a 2:1 dextran-to-PEG ratio, not to exceed 600 total μL in one 1.5 mL tube. This creates an aqueous two-phase system, where the dextran and the PEG solutions separate and are able to emulsify. The aqueous two-phase system was vortexed at full strength for 5 secs and immediately irradiated with 67.62 mW cm^-2^ of 320-390 nm UV light for 90 secs (Omnicure S1500, Excelitas Technologies). Microgel/dextran mixture was diluted in a solution of 0.1% w/v Pluronic F-127 (P2443, MilliporeSigma) in PS-PBS (Pluronic-PBS) until the microgels could be centrifuged down at 4000 xG for 3 mins. Supernatant was removed and microgels were resuspended in Pluronic-PBS and pelleted again at 2000 xG for 3 mins. Supernatant was removed and resuspended in Pluronic-PBS and passed through a 30 μm strainer (43-50030, pluriSelect) to be collected in a 50 mL conical. Filtered microgels were centrifuged at 2000 xG for 3 mins and supernatant removed and resuspended to < 1.5 mL with Pluronic-PBS to be able to fit in a 1.5 mL tube. The microgels were centrifuged at 2000 xG for 3 mins and all of the supernatant was removed. Microgel volume was measured and recorded by adding 400 μL of Pluronic-PBS and measuring the final volume of the mixture and subtracting 400 μL from this volume. Necessary amounts of microgel were collected and washed twice with PS-PBS to wash off the Pluronic. Microgels were then resuspended 5x in the medium of choice.

### Atomic Force Microscopy Sample Preparation and Measurement

Silicon wafers were etched functionalized by sonicating in 1 M NaOH (415413, MilliporeSigma for 1 hr and incubating them in 5% v/v (3-mercaptopropyl)trimethoxysilane (175617, MilliporeSigma) in ethanol for 30 mins. Microgels were resuspended in 1 mg/mL of PEG-dithiol in PBS and placed on top of the silicon wafer and allowed to settle for 5 mins before exposing to 67.62 mW cm^-2^ of UV light for 60 secs. Silicon wafers were washed three times with PBS. A Cypher AFM (Asylum Research) with biosphere AFM tips (NT_B2000_v0030, nanotools) was used to measure the microgels’ Young’s modulus. An area scan of 5 nN indentations was performed near the z-height top of the microgel and the top-most indentation was chosen for consideration. The indentation was fit with the Hertz model and a Poisson ratio of 0.5 was assumed.^[62,63]^

### Microgel Size Measurement

Microgels were placed on 8-well ibidi glass coverslips (80807, ibidi) and brightfield imaged using a widefield microscope (Axio Observer.Z1/7, Zeiss). Images were obtained at 20x and 0.8 NA. Microgel diameters were manually measured using ImageJ.

### Microwell Formation

8-well ibidi glass coverslips were plasma treated for 1 min and incubated with 2% v/v 3-(Trimethoxysilyl)propyl methacrylate (440159, MilliporeSigma) in ethanol for 15 mins, washed three times with ethanol and pressure air-dried. 2 μL of 10% w/v 4-arm PEG acrylate (10 kDa, 4arm-PEG-ACRL-10k, Laysan Bio) in 0.1% LAP in PS-PBS were dropped in each well and sandwiched with an 8 mm coverslip and irradiated with 67.62 mW cm^-2^ UV light for 30 secs. 8 mm coverslips were removed and 80 μL of PEG acrylate solution were added on top of the PEG puck and flattened with a PDMS mold with a 7 × 7 pillar array of 500 μm diameter and 200 μm pillars. The platform was irradiated with 67.62 mW cm^-2^ UV light for 30 secs and the mold removed to create a 7 × 7 array of 500 μm diameter and 200 μm wells.

### 2D Cell Culture

Tissue culture treated flasks were incubated for 30 mins in 1.07 µg cm^-2^ of poly-L-lysine (PLL, P6516, MilliporeSigma) at room temperature. Primary human hepatic stellate cells (HSCs) were sourced from Samsara Sciences and generously provided by Dr. Salman Khetani (University of Illinois Chicago) and cultured between passages 20-22 in the PLL-treated flasks at 37 °C and 5% CO_2_. DMEM with L-glutamine, 4.5 g L^-1^ glucose, sodium pyruvate and phenol red (10013CM, Corning) supplemented with 10% fetal bovine serum (FBS, 35010CV, Corning), 1% PenStrep and 1% L-glutamine (SH3003401, Corning) was used as culture medium during expansion (Exp. Media).

### 3D Cell Culture for Immunofluorescence Staining and RT-qPCR

25 µL of 5X microgel solution was mixed with 25 µL of 1.25 million HSCs per mL solution. This corresponds to an approximate volume ratio of 5:1 between pure microgels and pure cells. The volume corresponding to 2.5 μL of microgels (25 μL of microgel/cell solution) was dropped onto each microwell platform for immunostaining and 5 μL of microgels (50 μL of microgel/cell solution) were added onto three wells of an ultra-low attachment round bottom 384-well plate (10185-096, Corning) for RT-qPCR. All platforms were centrifuged at 300 xG for 1 min. RT-qPCR samples were placed in the incubator for 3 total days at 37 °C and 5% CO_2_. Immunostaining samples were washed three times with 400 μL of Exp. Media to remove all microgels and cells not captured within the microwells and placed in the incubator for 3 total days with a final Exp. Media volume of 250 μL at 37 °C and 5% CO_2_. RT-qPCR samples had half of its medium replaced after 1 day and immunostaining samples had all of its medium replaced after 1 day.

### Immunofluorescence Staining and Analysis

All washes were done carefully as to not disrupt the inside of the microwells. Microwells had their media removed and were fixed with 4% paraformaldehyde (RT15710, Electron Microscopy Sciences) in PBS for 20 mins. Microwells were then washed twice with PBS and permeabilized with 0.5% Triton X-100 (X100, MilliporeSigma) in PBS for 15 mins. Microwells were washed once with PBS and blocked with 1% w/v bovine serum albumin (BSA, A2153, MiliporeSigma) in 0.1% Triton X-100 in PBS for 1 hour. Microwells were washed once with PBS and stained for 24 hrs with 2 µg mL^-1^ of human pro collagen I alpha 1 antibody (AF6220, R&D Systems) and 0.848 µg mL^-1^ anti-LOX antibody (ab174316, abcam) or 4 µg mL^-1^ human PDGFR beta antibody (AF385, R&D Systems) and 10 µg mL^-1^ human/mouse/rat alpha-smooth muscle actin antibody (MAB1420, abcam) in 0.1% BSA and 0.1% Triton X-100 in PBS for 24 hrs in a gentle shaker. Microwells were washed three times with PBS with 15 min soaking intervals. Microwells were then stained for 4 hrs with 10 µg mL^-1^ of donkey anti-sheep IgG NL493-conjugated antibody (NL012, R&D Systems) and 10 µg mL^-1^ donkey anti-rabbit IgG Alexa Fluor 555-conjugated antibody (ab150062, abcam) or 10 µg mL^-1^ donkey anti-mouse IgG DyLight 550 (ab98795, abcam) and 10 µg mL^-1^ donkey anti-goat IgG DyLight 488 antibody (ab96935, abcam). Both antibody solutions were in 1 µg mL^-1^ DAPI (57-481-0, Fisher Scientific) and 200x DiD (V22889, ThermoFisher) in 0.1% BSA and 0.1% Triton X-100 in PBS. Microwells were washed three times with PBS with 15 mins soaking intervals and finally placed in 250 μL of 80% glycerol solution.

Samples were imaged using a Zeiss LSM 880 confocal laser scanning microscope with a 25x NA0.8 immersion objective and 1.5 RI oil at 30 °C. Z-stack images with a scale of 0.55 × 0.55 × 1.5 µm voxels were taken at a resolution of 1024 × 1024 per z slice. Images were analyzed using Imaris 9.7.0. Distance between nuclei was measured by segmenting the nuclei through Imaris, obtaining each nucleus’s x,y,z position and feeding these data points to a Python script that determines each nucleus’s shortest distance to another nucleus. These values were averaged within the biological replicate to produce one data point. Cellular structures were determined using the DiD stain and the total intracellular fluorescence intensity within the image was divided by the number of nuclei within the image to obtain the fluorescence intensity per nucleus. These values were normalized against the average fluorescence intensity per nucleus of all the image sets within the experimental replicate.

### RT-qPCR

HSCs from the three replicate wells were collected into one tube, centrifuged to 300 xG for 3 mins and dissociated using RLT buffer as recommended by RNeasy Plus Mini Kit (74134, Qiagen). All RNA isolation steps were done as recommended by the RNeasy kit. mRNA was quantified using a NanoDrop (ThermoScientific). mRNA was reverse transcribed as indicated by iScript Reverse Transcription Supermix (1708841, Bio-Rad). Amplification steps were performed in SsoAdanced Universal SYBR Green Supermix (1725272, Bio-Rad) using 250 nM of corresponding forward and reverse primers (Table 1). Reverse transcription and amplification were performed using a CFX Connect Real-Time PCR Detection System (Bio-Rad). Expression was measured as -ΔΔCT using GAPDH as the housekeeping gene.

**Table 1.**
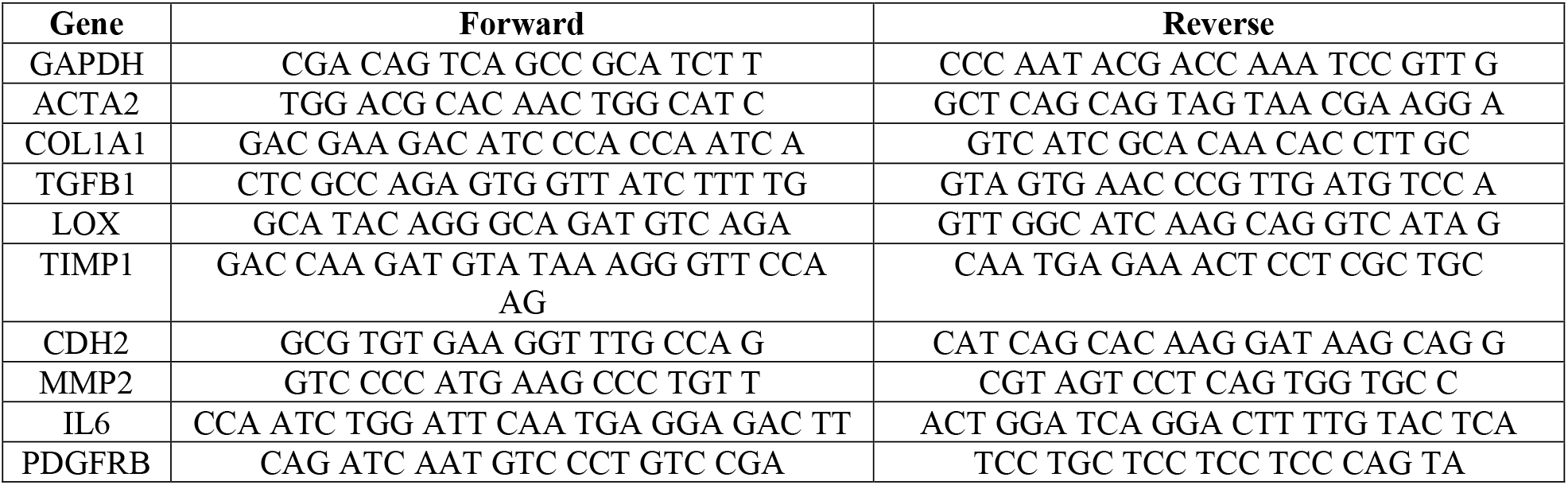
Forward and reverse primers used for the RT-qPCR reactions.

### High Throughput ECM Screening Platform and Plate Reading

4FN, 4C1, 4C3, 4C4, 8FN, 8C1, 8C3 and 8C4 microgels were prepared as indicated in PEG Microgel Formation and resuspended to 5 times the volume in DMEM with 4.5 g L^-1^ glucose, sodium pyruvate without L-glutamine and phenol red (17205CV, Corning) supplemented with 2% fetal bovine serum, 1% PenStrep and 1% L-glutamine (HTS Media). Microgels were distributed into 384-well black wall, flat, clear bottom plates (4588, Corning) as single component (24 µL), double component (12 + 12 µL) or triple component conditions (8 + 8 + 8 µL) using an OT-2 liquid handler (Opentrons) for a total of 4.8 µL of microgels each well. The plate was placed in an incubator overnight. 2D cultured HSCs were collected and 24 µL of 12.5 million cells per mL were manually distributed into each well and mixed thoroughly. Cells were cultured for 2 days in 37 °C and 5% CO_2_.

For MMP2 readouts, 24 µL of 37.5 µM MMP2 substrate (444212, MilliporeSigma) in 600 mM NaCl, 150 mM Tris-HCl, 15 mM CaCl_2_, 60 µM ZnSO_4_, 0.15% Brij-35 and 0.41% DMSO were added to each well for a total volume of 72 µL. Solutions were distributed rapidly using a 24-channel repeat pipettor. The plate was shortly centrifuged at 100 xG. After 6 hrs of incubation in 37 °C and 5% CO_2_, each well’s fluorescence was measured for emission at 383-403 nm wavelength, read height of 7 mm and at 37 °C with excitation at 315-335 nm wavelength using a Cytation5 (Agilent). All values were subtracted by a control with only 8FN microgels and no cells.

For metabolic readouts, 24 µL of 30% resazurin (AR002, R&D Systems) and 70% HTS Media was added onto each well of the plate for a total volume of 72 µL. Solutions were distributed rapidly using a 24-channel repeat pipettor. The plate was shortly centrifuged at 100 xG. After 4 hrs of incubation in 37 °C and 5% CO_2_, each well’s fluorescence was measured for emission at 580-600 nm wavelength, read height of 7 mm and at 37 °C with excitation at 534-554 nm wavelength using a Cytation5 (Agilent). All values were subtracted by a control with only 8FN microgels and no cells.

## ACKNOWLEDGEMENTS

The authors acknowledge the Institute for Genomic Biology for help and advice on imaging. The authors acknowledge the Materials Research Laboratory for help in manufacturing soft lithography materials and advice on atomic force microscopy. The authors thank Dr. Salman Khetani (UIC) for providing primary human hepatic stellate cells. Resazurin and resorufin chemical structure images obtained from Wikimedia commons.

## FUNDING SOURCES

This work was supported by NIH grant R01DK115747 (to GHU).

## Notes

### Competing Interest Statement

The authors have declared no competing interest.

