## Supplementary Information for "Combinatorial Microgels for 3D ECM Screening and Heterogeneous Microenvironmental Culture of Primary Human Hepatic Stellate Cells"

Supplementary Figure 1. Protein distribution in microgels depending on protein integration step.

Supplementary Figure 2. Immunofluorescence quantification for lysyl oxidase and α smooth muscle actin in HSCs cultured with microgels.

Supplementary Figure 3. RT-qPCR results for HSCs cultured in microgel scaffolds for ACTA2, CDH2, IL6, LOX, PDGFRB, and TIMP1.

Supplementary Figure 4. MMP2 and Resazurin readouts with 95% confidence intervals ranked based on average value.

Supplementary Figure 5. Linear regression coefficients for MMP2 and resazurin readouts of subset data for 2 and 3 component conditions.

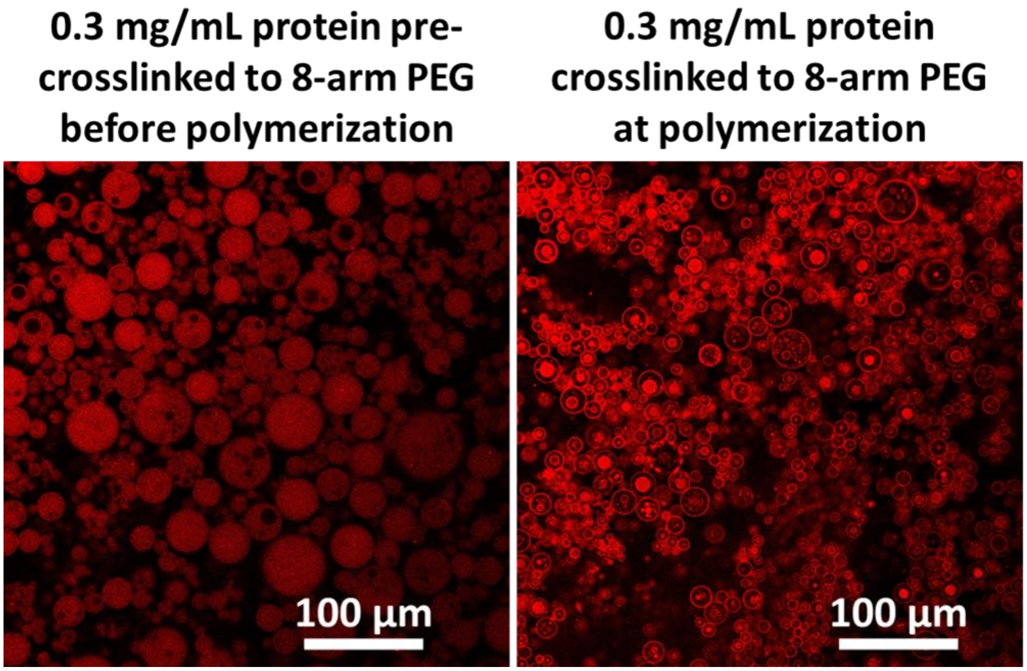


**Supplementary Figure 1.** Distribution of SVA-PEG-SH functionalized Alexa Fluor 555 tagged antibody in 8-arm 8% PEG norbornene microgels at 0.3 mg $mL^{-1}$ protein and 100 µM SVA-PEG-SH final concentration. When the antibody was pre-crosslinked to the PEG norbornene, the antibody was more evenly distributed across the microgel. When the antibody was crosslinked to the PEG norbornene at the time of polymerization, the antibody tended to aggregate on the outside or core of the microgel. Red: Alexa Fluor 555 antibody.

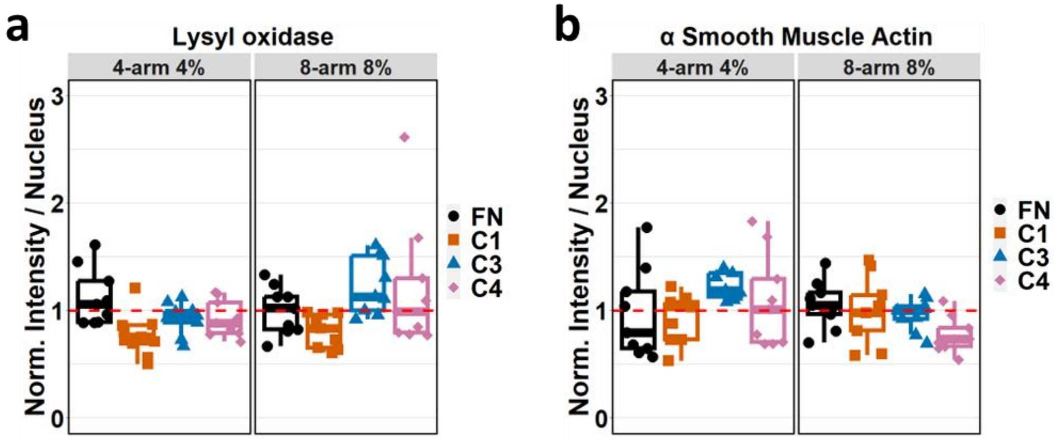


**Supplementary Figure 2.** Immunofluorescence quantification for HSCs cultured with microgels. **a)** Box and whisker plots of the anti-lysyl oxidase fluorescence intensity per nucleus normalized to the average of all conditions per experimental replicate. n = 9 from 3 experimental replicates. **b)** Box and whisker plots of the anti-αSMA fluorescence intensity per nucleus normalized to the average of all conditions per experimental replicate. n = 9 from 3 experimental replicates. No significant differences were observed between any of the conditions within the same PEG norbornene composition or same ECM protein component.

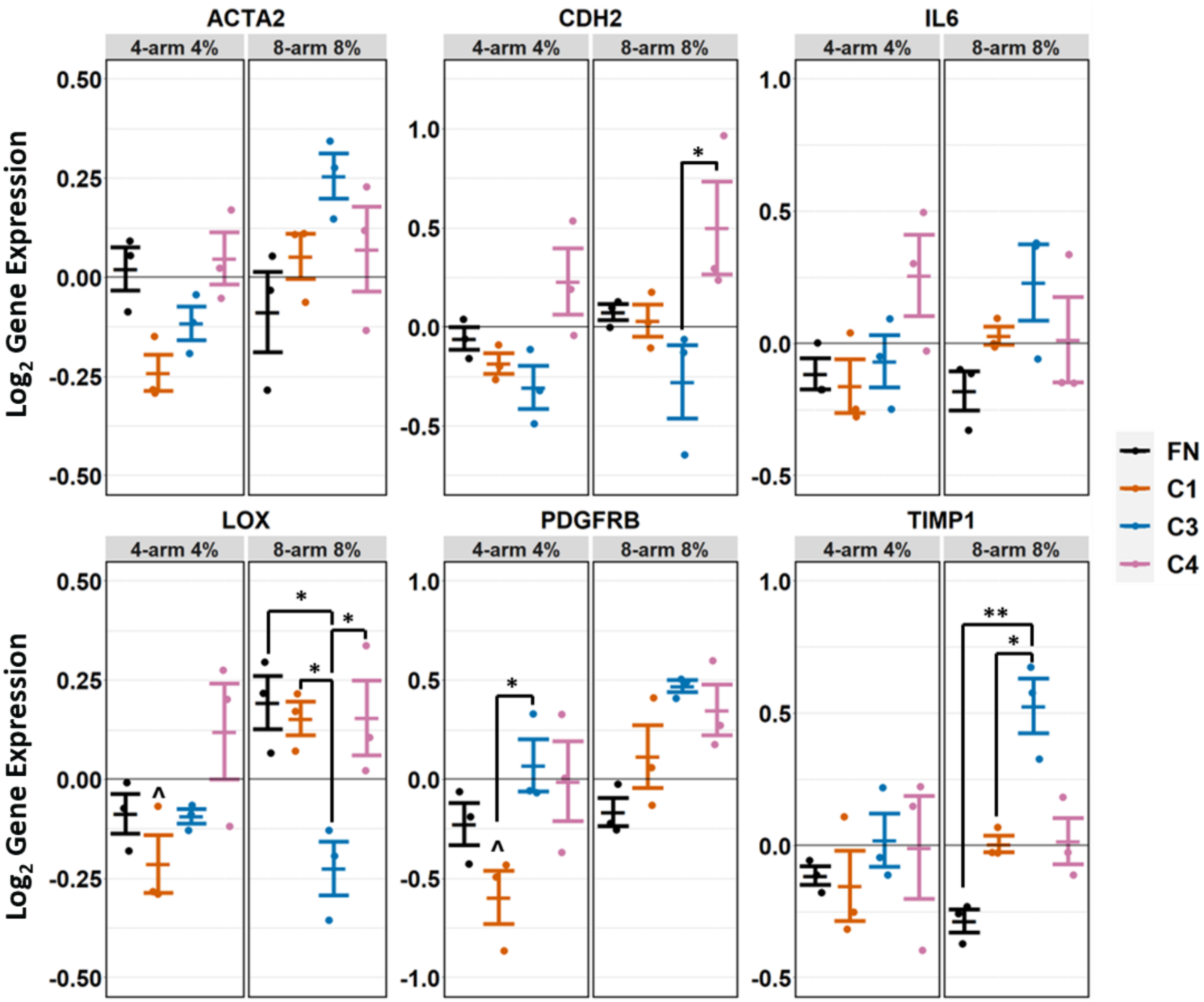


**Supplementary Figure 3.** Average ± standard error of means for the genes not shown individually in Figure 4. n = 3 experimental replicates. Two-way interaction ANOVA analysis. * $p < 0.05$ ** $p < 0.01$ where * means between conditions and ^ against its 8-arm 8% counterpart.

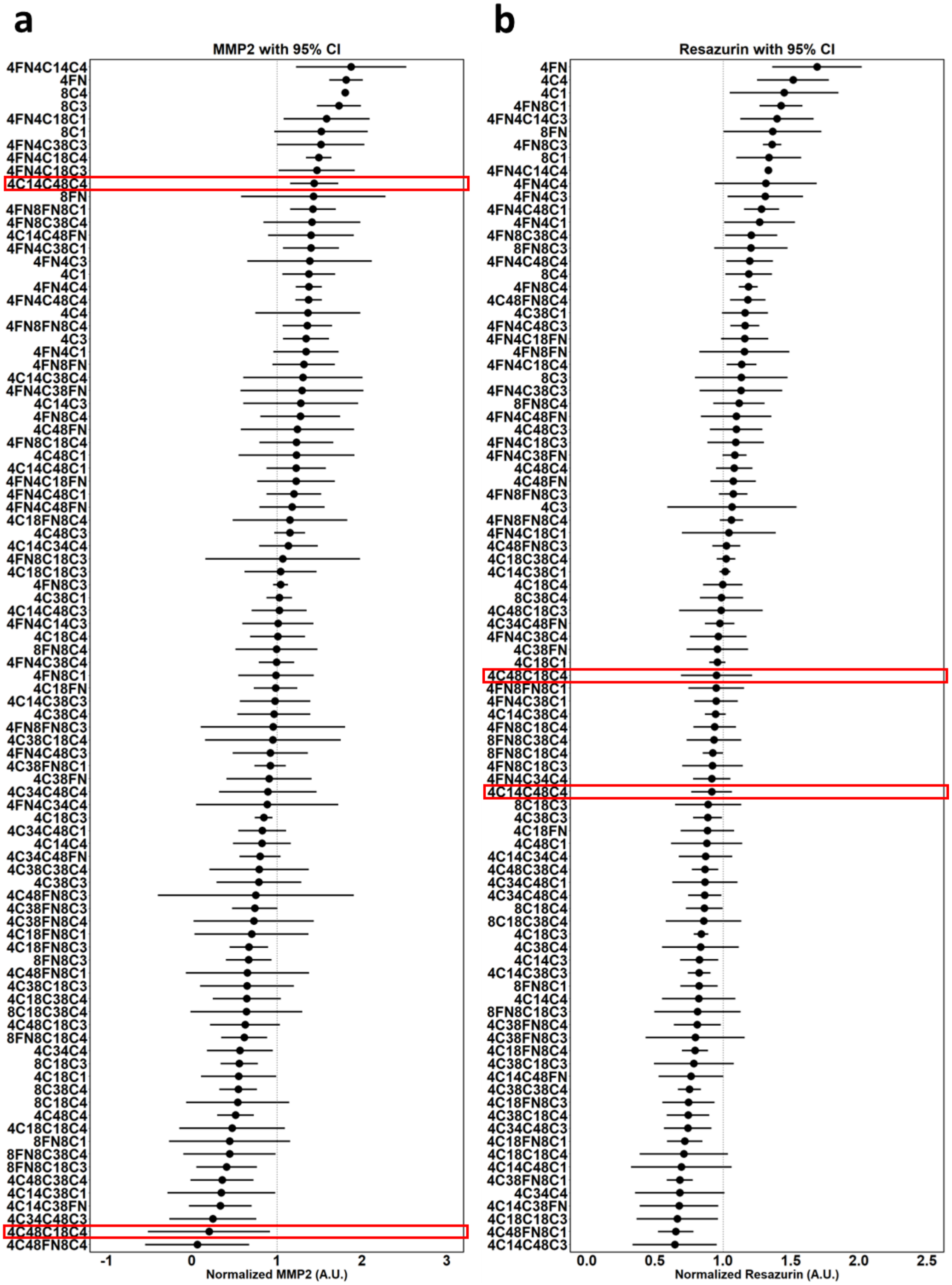


**Supplementary Figure 4.** Normalized plate reading readouts for all the conditions tested with their corresponding 95% confidence intervals. **a)** MMP2 substrate readout. **b)** Resazurin readout.

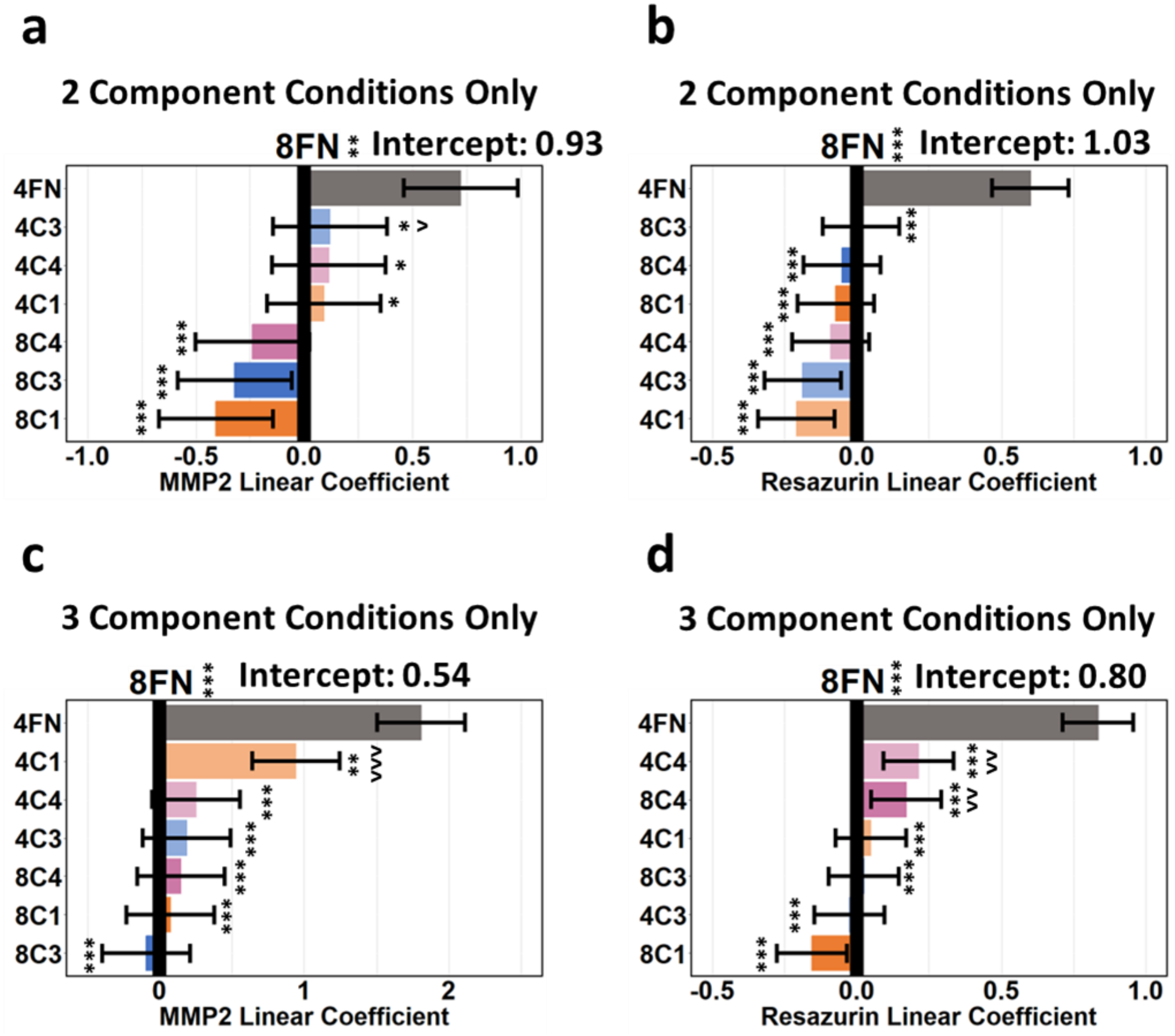


**Supplementary Figure 5.** Linear regression analysis performed on subsets of plate readout data. **a)** Linear coefficients of MMP2 readout for only 2 component conditions. **b)** Linear coefficients of resazurin readout for only 2 component conditions. **c)** Linear coefficients of MMP2 readout for only 3 component conditions. **d)** Linear coefficients of resazurin readout for only 3 component conditions.
